# Heterozygous *Eif4nif1* Stop Gain Mice Replicate the Primary Ovarian Insufficiency Phenotype in Women

**DOI:** 10.1101/2024.04.09.588694

**Authors:** Mika Moriwaki, Lihua Liu, Emma R James, Neal Tolley, Ashley M O’Connora, Benjamin Emery, Kenneth Ivan Aston, Robert A. Campbell, Corrine K. Welt

**Affiliations:** Division of Endocrinology, Metabolism and Diabetes, University of Utah School of Medicine, Salt Lake City, UT, USA; Department of Cell Biology & Physiology, University of New Mexico Health Sciences Center, Albuquerque, NM, USA; Andrology and IVF Laboratory, Department of Surgery (Urology), University of Utah School of Medicine, Salt Lake City, UT, USA; Program in Molecular Medicine, University of Utah, Salt Lake City, UT, USA; Biomedical Informatics Core of the Clinical and Translational Science Institute, University of Utah, Salt Lake City, UT, USA; Department of Internal Medicine, University of Utah, Salt Lake City, UT, USA; Division of Microbiology and Immunology, Department of Pathology, University of Utah, Salt Lake, City, USA

**Keywords:** Key terms: reproduction, fertility, embryogenesisd

## Abstract

We created the c.1286C>G stop-gain mutation found in a family with primary ovarian insufficiency (POI) at age 30 years. The *Eif4enif1* C57/Bl6 transgenic mouse model contained a floxed exon 10-19 cassette with a conditional knock-in cassette containing the c.1286C>G stop-gain mutation in exon 10. The hybrid offspring of CMV-*Cre* mice with *Eif4enif1^WT/flx^* mice were designated *Eif4enif1^WT/^*^Δ^ for simplicity. A subset of female heterozygotes (*Eif4enif1^WT/^*^Δ^) had no litters. In those with litters, the final litter was earlier (5.4±2.6 vs. 10.5±0.7 months; p=0.02). Heterozygous breeding pair (*Eif4enif1^WT/^*^Δ^ *x Eif4enif1^WT/^*^Δ^) litter size was 60% of WT litter size (3.9±2.0 vs. 6.5±3.0 pups/litter; *p*<0.001). The genotypes were 35% *Eif4enif1^WT/flx^* and 65% *Eif4enif1^WT/^*^Δ^, with no homozygotes. Homozygote embryos did not develop beyond the 4-8 cell stage. The number of follicles in ovaries from *Eif4enif1^WT/^*^Δ^ mice was lower starting at the primordial (499±290 vs. 1445±381) and primary follicle stage (1069±346 vs. 1450±193) on day 10 (p<0.05). The preantral follicle number was lower starting on day 21 (213±86 vs. 522±227; p<0.01). Examination of ribosome protected mRNAs (RPR) demonstrated altered mRNA expression. The *Eif4enif1* stop-gain mice replicate the POI phenotype in women. The unique mouse model provides a platform to study regulation of protein translation across oocyte and embryo development in mammals.

## Introduction

Primary ovarian insufficiency (POI) reflects a continuum from infertility in women with ovarian dysfunction to early menopause (1). Data support a genetic cause in women with POI (2–4). Using whole exome sequencing, we identified dominant inheritance of a stop-gain mutation (c.1286C>G, p.Ser429Ter) in *eIF4ENIF1* in a large family in which female carriers of the mutation developed menopause at approximately age 30 years (5). The affected women had no additional medical problems and carrier males had normal fertility, suggesting that the protein is ovary specific.

*eIF4ENIF1* codes for one of the binding proteins critical for repressing *eIF4E,* the rate limiting, 5’ cap-dependent translation initiation factor (6). As demonstrated in cell models, *eIF4ENIF1* regulates eIF4E translation initiation by binding at the same site used by the eIF4G scaffolding protein to bind to eIF4E, taking it out of its active complex (6,7)(Supplementary Figure 1). In the oocytes of non-mammalian species, *eIF4ENIF1* homologues target bound mRNA to localized sites in the oocyte or germ cell granules, where maternal mRNAs are stored for later translation during meiosis and embryogenesis (8–11). However, there is very little known about *eIF4ENIF1* and its role in mammalian oocytes.

We created a mouse model with the precise *eIF4ENIF1* stop gain mutation found in humans to determine the mechanism causing POI. The heterozygous *Eif4enif1* stop-gain mouse model replicated the decreased reproductive lifespan and early oocyte loss of human POI. The model also provides insight into the role of *eIF4ENIF1* during oocyte and embryo development.

## Methods

### Animals

The *Eif4enif1* C57/Bl6 transgenic mouse model (KOMP, Davis, CA) contains a floxed exon 10-19 cassette and a conditional knock-in cassette containing exon 10 with the c.1283C>G stop-gain mutation, recreating the mutation causing familial POI in the mouse, and WT exons 11-19 (*Eif4enif1^WT/flx^*, Figure 1) (5). Both cassettes contain the WT STOP codon, polyA and 3’UTR. All mice were the hybrid offspring of CMV-*Cre* mice with *Eif4enif1^WT/flx^*mice and designated *Eif4enif1^WT/^*^Δ^ for simplicity. Genotype analyses were performed by PCR with DNA extracted from ear or tail biopsies and Sanger sequencing. Primers used to detect WT cDNA and mutant cDNA in the knock-in construct were 5’-TACAGCAGGGTTGTAGCAGC-3’ and 5’-CACTGTGGTCTCAAGGCAC-3’. Primers used to detect a stop-gain mutation in *Eif4enif1* were 5’-TTCTAAACACCCTTTGCTTGG-3’ and 5’-CCCTTCAGCCCTGCTTCTAC-3’. Primers to detect the *Cre* transgene alleles were 5’-TCTCACGTACTGACGGTGG-3’ and 5’-ACCAGCTTGCATGATCTCC-3’.

**Figure 1.**
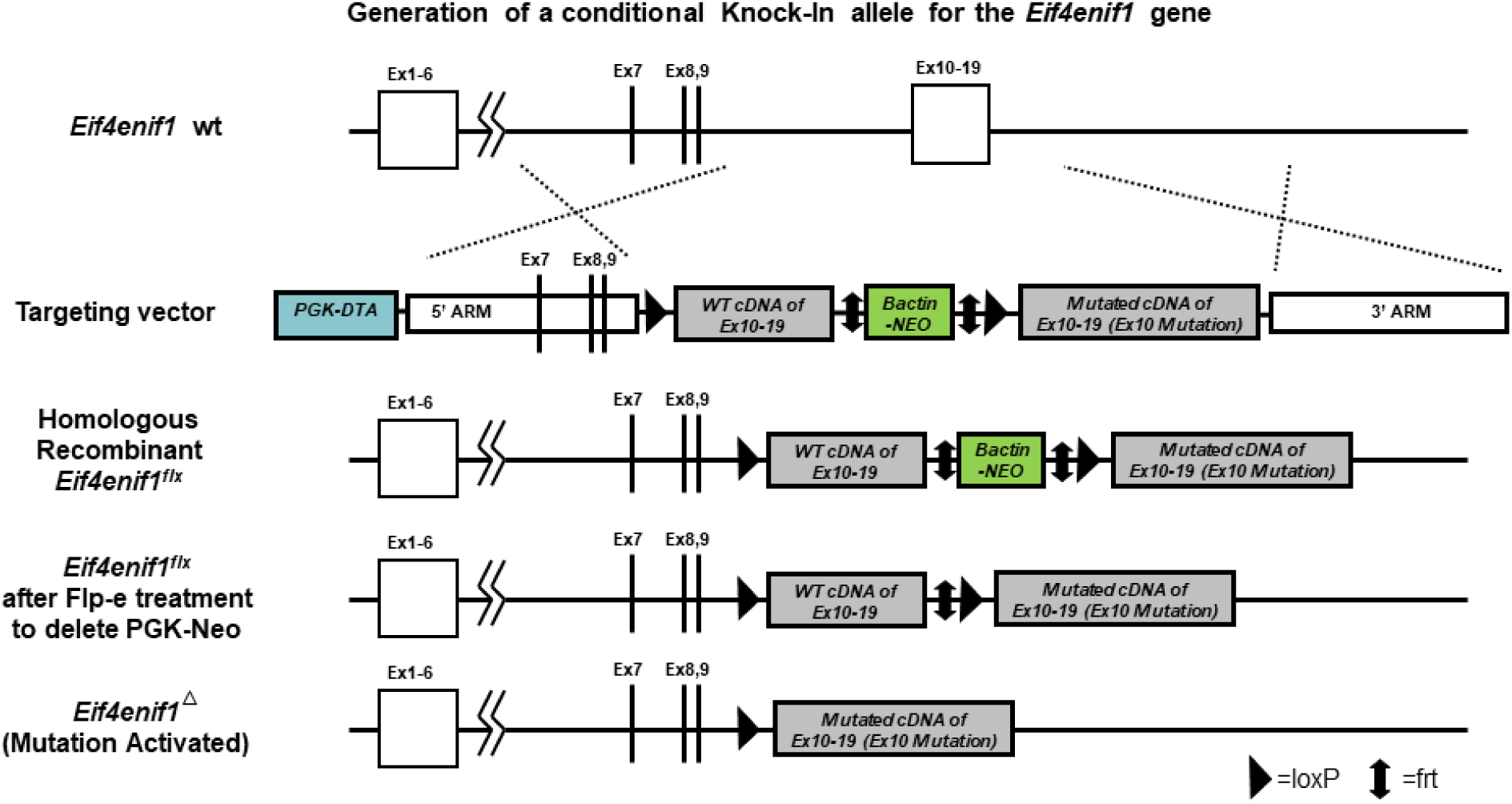
Generation of a conditional Knock-In allele for the Eif4enif1 gene. The Knock-In cassette (grey) consists of LoxP sites that flank a “WT cDNA insert” and an frt-flanked Neomycin (Neo) marker (green). This is followed by a “Mutant cDNA insert” to complete the Knock-In cassette (grey). This Knock-In cassette replaced the endogenous exons10-19 through homologous recombination. In the targeting vector, this Knock-In cassette is flanked by arms of homology. The 5’arm is 9.3kb and contains exons 7-9, while the 3’arm is 9.4kb and targets just downstream of the Eif4enif1 gene. The construct also contains a Diphtheria Toxin A (DTA) cassette (blue). The Neo element allowed for positive selection in ES cells, while the DTA element permitted negative selection in ES cells. The “WT cDNA insert” was comprised of WT cDNA from exons10-19, and thus contains the STOP codon, polyA and 3’UTR. The “Mutant cDNA insert” is made up of Mutated cDNA from exons10-19, and also contains the STOP codon, polyA and 3’UTR. The sole mutation in the Mutant cDNA is in base pair #7 of exon10 where C is changed to G, which results in the serine residue being converted to a STOP codon. After homologous recombination of the conditional knock-in construct, the Neo was excised via Flp-e administration. The resulting Eif4enif1 gene had normal expression until Cre-mediated deletion of the WT cDNA insert. This deletion conditionally activated the mutation contained in the Mutant cDNA, which resulted in a truncated transcript and protein.

All experimental animal procedures were approved by the Institutional Animal Care and Use Committee at the University of Utah. Animals were obtained from the Transgenic Gene-Targeting Mouse Facility at the University of Utah (Salt Lake City, UT) and housed in a pathogen-free environment with controlled temperature, humidity, and 12-hour light/dark cycle.

### Tissue Preparation for Histological Analyses

Phenotypic characteristics, body weights, litter counts and timing were recorded. Mice younger than 21 days of age were euthanized by isoflurane (VetOne, Boise, Idaho) followed by cervical dislocation. Mice older than 21 days of age were euthanized by CO_2_ asphyxiation.

Ovaries were collected from mice at each target age. Tissues were stored in 70% ethanol and subsequently fixed in paraformaldehyde, embedded in paraffin, sectioned to a thickness of 6 μm, and affixed on positively charged glass slides. Oven-dried sections were deparaffinized in xylene, rehydrated by a graded ethanol series (100%, 95%, 70%, 50%, and 0%), and processed for further histological analyses.

### Follicle Counts

Follicles were counted in fixed ovaries from mice aged days 1-5 (primordial and primary follicles), day 7, day 10, day 22 (first wave of growing follicles from small preantral to small antral follicles), week 20 (peak fertility), then every 2 months from 10 months to 26 months (follicle exhaustion)(12–14). Hematoxylin and eosin (H &E) staining was performed using an H&E staining kit (BBC Biological, Mount Vernon, VA). For all sections, primordial and primary follicles were counted in every fourth section (6 µm sections X 3 sections = 18 µm) to avoid double counting follicles with a mean follicle diameter of < 20 μm. All preantral and antral follicles were counted only once, in the plane of the nucleus. Follicles were divided into four groups: 1) primordial, 2) primary, 3) preantral, and 4) antral follicles; along with 5) corpora lutei. Data are analyzed using two-way analysis of variance (ANOVA) with post hoc testing.

### Immunofluorescence

After deparaffinization and rehydration, slides went through the same antigen retrieval, blocking, and primary antibody incubation as for immunohistochemistry. Primary antibodies used were rabbit C-terminal eIF4ENIF1 (1:10,000, A300-706A, Bethyl, Montgomery, TN), mouse eIF4E (1:100, MA1-089, Thermo Fisher Scientific, Waltham, MA), and rabbit DDX6 (1:200, PA5-27786, Thermo Fisher Scientific, Waltham, MA). A second set of ovaries was imaged using N-terminal eIF4ENIF1 rabbit antibody for comparison (2 μg/ml, NBP1-89389, Novus Biologicals, Littleton, CO). Slides were incubated in anti-mouse Alexa Fluor 488 and anti-rabbit Alexa Fluor 594 (Thermo Fisher Scientific, Waltham, MA) diluted in 1× TBS with 1% BSA for 1 hour at RT in a dark room. Slides were counterstained with DAPI (SouthernBiotech, Birmingham, AL) and mounted with glycerol mounting medium with DABCO (Electron Microscopy Sciences, Hatfield, PA). Images were captured on the Nikon fluorescent microscope and NIS Elements software (Nikon Instruments Inc., Melville, NY).

Oocytes and fertilized embryos at varying stages were also incubated with rabbit C-terminal eIF4ENIF1 (1:10,000, A300-706A, Bethyl, Montgomery, TN) or rabbit DDX6 (1:200, PA5-27786), then incubated in anti-rabbit Alexa Fluor 594 (Thermo Fisher Scientific), as above, and counterstained with DAPI (SouthernBiotech).

### Immunohistochemistry (IHC)

IHC was performed on day 5, 17, 22, week 20 and year 1. Rehydrated slides were submerged in sodium citrate buffer (10 mM sodium citrate, 0.05% Tween-20, pH 6.0) at 95[ for 20 minutes for antigen retrieval, washed in TBS with 0.025% Triton X-100 (TBST), blocked in 10% donkey normal serum (Abcam, Cambridge, MA) for 2 hours at room temperature (RT), and incubated with rabbit active Caspase-3 (5 μg/ml, AF835-SP, R&D systems, Inc., Minneapolis, MN), rabbit cleaved Caspase-9 (1:50, 9509, Cell Signaling Technology, Danvers, MA), and rabbit cleaved PARP (1:100, 94885, Cell Signaling Technology, Danvers, MA) diluted in TBS with 1% BSA at 4[ overnight. Endogenous peroxidase activity was suppressed in 0.3% H2O2 in 1× TBS for 15 minutes. Slides were incubated in HRP-conjugated anti-rabbit secondary antibody (Thermo Fisher Scientific, Waltham, MA) diluted in TBS with 1% BSA for 1 hour at RT, rinsed with TBS, developed with 3, 3’-Diaminobenzidine (DAB, Acros Organics, Pittsburgh, PA), washed with water, counterstained with hematoxylin, dehydrated, and mounted with Permount mounting medium (Fisher Scientific, Waltham, MA). The Apoptag Plus Peroxidase *In Situ* Apoptosis Detection Kit (MilliporeSigma, Burlington, MA) was used to perform the TUNEL assay according to the manufacturer’s directions. The compound light microscope (Carl-Zeiss, Oberkochen, Germany) was used for image acquisition.

### Superovulation & IVF

Week 7-10 *Eif4enif1 ^WT/^*^Δ^ and *Eif4enif1 ^WT/flx^* mice underwent intraperitoneal injection with 5 IU PMSG to initiate follicular development and 5 IU hCG 48 hours later to induce ovulation (15). Sperm were collected under oil from the caudal epididymis and vas deferens week 9-10 *Eif4enif1 ^WT/^*^Δ^ or *Eif4enif1 ^WT/flx^* males using a standard swim out approach. After semen analysis, oocytes were fertilized by *in vitro* fertilization (IVF). Sperm was collected and placed in fertilization medium for 45 min to stimulate capacitation prior to harvesting oocytes from super-ovulated females. Thirteen hours after hCG administration, females were euthanized and oviducts removed. Each oviduct was placed in a wash drop and dissected to obtain cumulus oocytes complexes. Following enzymatic removal of cumulus cells with hyaluronidase, metaphase II oocytes were transferred to 100 µl fertilization drops containing ∼2.5 x 10^6^ capacitated, motile sperm. After four hours oocytes were transferred to KSOM-AA drops, and development was assessed from pronuclear to the blastocyst stage.

Embryos were categorized in developmental stages including unfertilized, pronuclear, two-cell, four-cell, eight-cell, morula and blastocyst stages with an additional categorization for unhealthy, dead or fragmented embryos. Each embryo produced by IVF was lysed in 50 μl digestion buffer comprising 10 mM Tris HCl at pH 8.3, 50 mM KCl, 0.1 mg/ml gelatin, 0.45% NP40, 0.45% Tween-20, and 200 μg/ml proteinase K at 55[ for 5 minutes (16). Individual embryos were genotyped for *Eif4enif1* alleles as described above.

### Western Blot

Snap-frozen ovaries at day 1, 5 and 10 were lysed in ice-cold RIPA buffer consisting of 50 mM Tris HCl (pH 8.0), 150 mM NaCl, 1% NP-40, and 0.5% SDS, and protease inhibitor (Fisher Scientific, Waltham, MA) for 30 minutes at 4[ and cell debris was removed by 12,000 rpm centrifugation for 20 minutes at 4[. Protein concentrations of the resulting lysates were assessed using the DC protein assay (Bio-Rad, Hercules, CA). Proteins in cell lysates were heated at 95[ for 5 minutes in Laemmli samples buffer (Bio-Rad, Hercules, CA), resolved by 10% SDS-PAGE, and transferred onto nitrocellulose membranes. The membranes were blocked in 5% non-fat milk in TBST (50 mM Tris-HCl and 150 mM NaCl with 0.1% Tween-20 adjusted at pH 7.6) for 1 hour at RT and incubated in primary antibodies diluted in 2% non-fat milk in TBST at 4[ overnight. The primary antibodies used were eIF4E rabbit antibody (1:1,000, PA5-24582, Thermo Fisher Scientific), C-terminal eIF4ENIF1 rabbit antibody (1:10,000, A300-706A), N-terminal eIF4ENIF1 rabbit antibody (1:1000, NBP1-89389, Novus Biologicals, Littleton, CO) and β-actin rabbit antibody (1:5000, NB600-532SS, Novus Biologicals, Littleton, CO). The membranes were incubated with an anti-rabbit HRP-conjugated secondary antibody (Thermo Fisher Scientific, Waltham, MA) for 1 hour at RT. The signals were developed by ECL solutions and detected by standard X-ray films (Thermo Fisher Scientific, Waltham, MA). Band density was analyzed using VisionWorks LS software (UVP, Upland, CA). All experiments were repeated three times. Relative Eif4enif1 quantities measured using N and C-terminal antibodies were controlled for Bactin and compared using one-way ANOVA within a time point and two-way ANOVA to examine differences across time, with Holm-Sidak post hoc testing. A *p* value <0.05 was considered significant.

### RNA Seq and Ribosome Footprint Profiling

Methods have been previously published (17,18). For Ribo-Seq, ovaries each from day 5 (n=5 each), day 10 (n=4 each) and day 22 (n=2 each) *Eif4enif1 ^WT/^*^Δ^ and *Eif4enif1^WT/flx^* mice were prepared. Ovaries were removed, placed into a buffer containing cycloheximide, shredded with a needle, frozen in liquid nitrogen and stored at −80°C. Cell lysates were later thawed to room temperature, placed into lysis buffer and homogenized.

Total RNA was extracted from homogenized samples using the RNeasy mini kit and treated with DNaseI (QIAGEN, Hilden, Germany). RNA-seq libraries were prepared with NEBNext Ultra I Directional RNA Library Prep with rRNA Depletion (New England Biolabs, Inc., Ipswich, MA). Reads were aligned (Novoalign) to the reference genome (M_musculus_Jun_2020, GRCm39) and a pseudotranscriptome containing splice junctions.

The Deseq2 analysis package was used to determine differential expression and significance according to the Deseq2 algorithm, which assigns reads to composite transcripts (1 per gene) and quantifies fragments per kilobase of transcript per million mapped reads (FPKMs). Multiple comparisons were controlled using a false discovery rate (FDR) according to the method by Benjamini-Hochberg. For differential expression of total (Tot) mRNAs, reads mapping to ribosomal RNA were removed. We considered genes that had at least 10 transcripts per million and a difference in expressed transcripts with a FDR <0.05 and a log_2_ fold change ≥ 2 for examination in *Eif4enif1 ^WT/^*^Δ^ vs. *Eif4enif1^WT/flx^* mice.

For ribosome footprint profiling, RNAse 1 was added to the tissue lysate and incubated for 45 min. The reaction was stopped with 10 µL Superase*IN range inhibitor and transferred to ice. The preparation was subsequently run on GES400 exclusion columns and monosome fraction collected. RNA was extracted from total input and fractionated output of polysome profiling using RNeasy mini kit and cleaned up with DNaseI (QIAGEN). RNA was purified on a 15% polyacrylamide gel and extracted from the gel. Ribosomal RNA was removed using the NEBNext rRNA depletion solution and Probe Hybridization buffer. DNAse1 was used to digest DNAse in the mixture. The ribosome profiling library was prepared using NEBNext Multiplex Small RNA Library Prep Set (New England Biolabs).

Differential expression was performed on ribosome protected mRNAs (RPRs). We considered genes that had at least 10 transcripts per million and a difference in expressed transcripts with a with a FDR <0.05 and a log_2_ fold change ≥2 for further analysis in *Eif4enif1 ^WT/^*^Δ^ vs. *Eif4enif1^WT/flx^* mice. Translational efficiency (TE) was calculated by subtracting log2 WT RPR from WT total and the log2 Het RPR from Het total.

### Polysome Profiling

Collagenase-treated ovaries were lysed by 3-minute incubation on ice with 375 μl low salt buffer supplemented with 30 μl/ml RNase inhibitor and 125 μg/ml cyclohexamide. The cell lysate was further lysed by 5-minute incubation on ice with 125 μl high salt lysis buffer and centrifuged at 20,800 ×g for 1 minute at 4[. The supernatant was loaded onto 10-50% sucrose gradients and centrifuged at 47,000 rpm for 1 hour at 4[ using SW55 Ti roter. Sucrose gradient fractions were monitored at 254 nm absorbance on an UA-6 UV/VIS detector (Teledyne Isco Inc., Lincoln, NE) and collected in individual tubes containing TRIzol LS (Ambion, Austin, TX).

RNA from polysome fractions of day 5 mouse ovaries was reverse transcribed to cDNA using the SuperScript VILO master mix (Thermo Fisher Scientific, Waltham, MA). To confirm the presence of the stop-gain mutation in actively translated mRNA, cDNA from polysome fractions of mouse ovaries was PCR amplified with *Eif4enif1* exon 9_10 primers (forward 5’-CCTCTTCTTTCTAGCCTTTCTGC-3’ and reverse 5’-TTCTGCCATGAAGGGAGTTG-3’) and genotyped by Sanger sequencing.

## Results

### Mouse Phenotype

The heterozygotes (*Eif4enif1^WT/^*^Δ^ ) have no outward phenotypic differences compared to WT (*Eif4enif1^WT/flx^*). Body weight was similar in *Eif4enif1^WT/^*^Δ^ vs. *Eif4enif1^WT/flx^*(two-way ANOVA female p=0.4, male p=0.2; Supplementary Figure 2). The lifespan of the *Eif4enif1^WT/^*^Δ^ and *Eif4enif1^WT/flx^* animals was similar (649±140 vs. 630±106 days; p=0.63).

### Gonad Size

There was no difference in testis volume / mass in male *Eif4enif1^WT/^*^Δ^ mice compared to *Eif4enif1^WT/flx^*(5.6±0.4 vs. 6.1±0.4 mm^3^/g; p=0.4; Supplementary Figure 3A). There was also no difference in ovarian volume between *Eif4enif1^WT/^*^Δ^ and *Eif4enif1^WT/flx^* (4.4±1.9 vs. 4.7±0.7 mm^3^; p=0.8; Supplementary Figure 3B).

### Mouse Fertility

Fertility was assessed by litter timing, litter counts and reproductive lifespan. The heterozygotes (*Eif4enif1^WT/^*^Δ^) have decreased fertility compared to control females (*Eif4enif1^WT/flx^*) when mated with control males (*Eif4enif1^WT/flx^*) or heterozygous males (*Eif4enif1^WT/^*^Δ^), as assessed by litter count and reproductive lifespan, but not litter interval (Table 1).

**Table 1.**
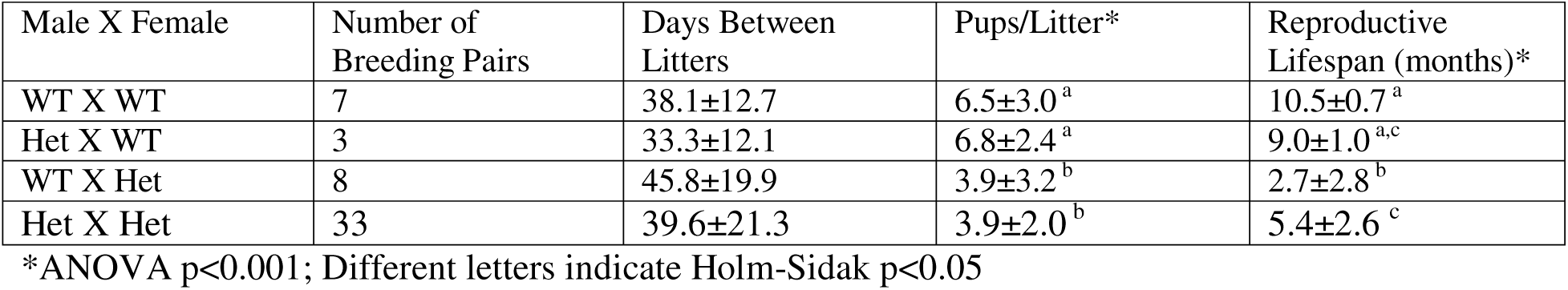
Fertility Information from *Eif4enif1^WT/flx^* (WT) and *Eif4enif1^WT/^*^Δ^ (Het) crosses.

| Male X Female | Number of Breeding Pairs | Days Between Litters | Pups/Litter* | Reproductive Lifespan (months)* |
| --- | --- | --- | --- | --- |
| WT X WT | 7 | 38.1±12.7 | 6.5±3.0<sup>a</sup> | 10.5±0.7<sup>a</sup> |
| Het X WT | 3 | 33.3±12.1 | 6.8±2.4<sup>a</sup> | 9.0±1.0<sup>a,c</sup> |
| WT X Het | 8 | 45.8±19.9 | 3.9±3.2<sup>b</sup> | 2.7±2.8<sup>b</sup> |
| Het X Het | 33 | 39.6±21.3 | 3.9±2.0<sup>b</sup> | 5.4±2.6<sup>c</sup> |

\*ANOVA p<0.001; Different letters indicate Holm-Sidak p<0.05

A subset of female heterozygotes (*Eif4enif1^WT/^*^Δ^) had no litters in 20 weeks (7 of 43; 16%). In those *Eif4enif1^WT/^*^Δ^ females with litters, the average length of time between litters was not different than for *Eif4enif1^WT/flx^* males and females or *Eif4enif1^WT/^*^Δ^ males with *Eif4enif1^WT/flx^* females (p=0.2; Table 1). The final litter was earlier in female *Eif4enif1^WT/^*^Δ^ mice (overall p<0.001).

Heterozygous breeding pair (*Eif4enif1^WT/^*^Δ^ *x Eif4enif1^WT/^*^Δ^) litter size was 63% of WT litter size (4.1±2.0 vs. 6.5±3.0 pups/litter; 0<0.001)(Table 1). The genotypes were 35% *Eif4enif1^WT/flx^* and 65% *Eif4enif1^WT/^*^Δ^, with no homozygotes.

### Follicle Counts

Follicle number was lower at all timepoints in *Eif4enif1^WT/^*^Δ^ compared to *Eif4enif1^WT/flx^* starting after day 5. Primordial follicles were decreased starting on day 10, primary follicles were decreased through month 12 and preantral follicles decreased through month 18 (Figure 2). The number of antral follicles was also lower but post hoc testing did not indicate differences on a specific day and there was no difference in the number of corpora lutei.

**Figure 2.**
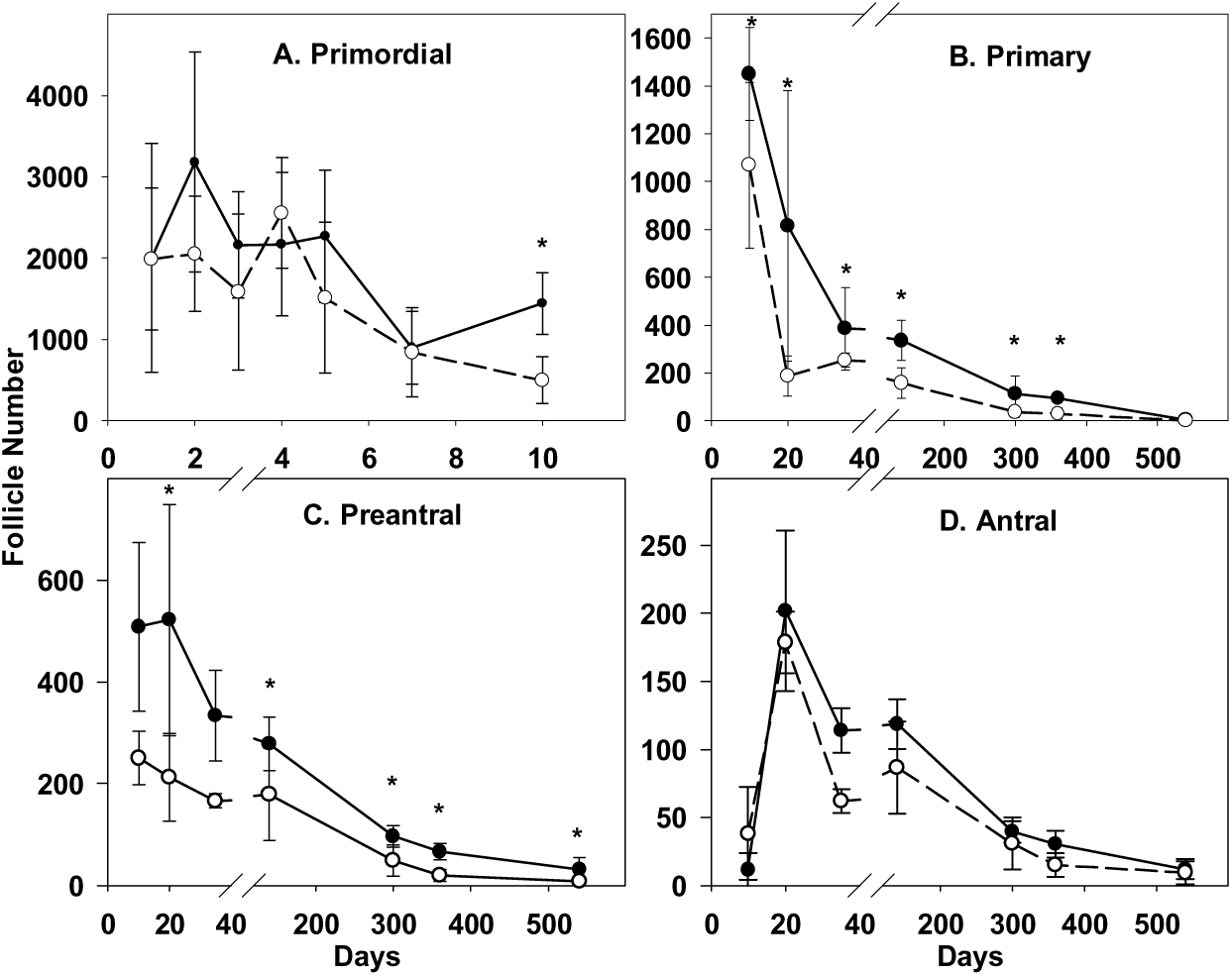
A) Primordial, B) primary, C) preantral and D) antral follicle number in *Eif4enif1^WT/flx^*(closed circles) compared to *Eif4enif1^WT/^*^Δ^ (open circles). Two way *ANOVA p*<0.01 day 10 through 18 months for primordial, primary and preantral follicles. *indicates Holm-Sidak post hoc test p<0.05.

### Oocyte Apoptosis

Caspase 3 and PARP were identified in the oocyte in a small number of preantral and small antral follicles in the *Eif4enif1 ^WT/^*^Δ^ but not *Eif4enif1^WT/flx^*oocytes on week 20 (Supplementary Figure 4) and caspase 3 staining was demonstrated in atrophic oocytes without clear surrounding granulosa cells at year 1. In addition, TUNEL staining on day 17 and day 22 demonstrated 10-28 positive granulosa cells in follicles in which the oocyte was pulled away from the edges, and up to 50% of the granulosa cells surrounding atrophic or missing oocytes in the periphery of the ovary of *Eif4enif1 ^WT/^*^Δ^ mice. In contrast, only 1-2 granulosa cells demonstrated staining in *Eif4enif1^WT/flx^* mice. There was no apparent caspase 3, PARP or TUNEL staining on day 5.

### Eif4enif1 Localization and Quantification

#### Immunofluorescence

We studied protein expression in ovaries from *Eif4enif1 ^WT/^*^Δ^ and *Eif4enif1^WT/flx^* mice on days 3, 10 (Figure 3 and 4) and 21 (not shown) and week 20 (Figure 5). Eif4enif1 immunofluorescence was present in oocyte and granulosa cell cytoplasm. Eif4enif1 immunofluorescence was less intense in *Eif4enif1 ^WT/^*^Δ^ than *Eif4enif1^WT/flx^* on days 3, 10 and 21 and on week 20 (Figure 3-5). There was no apparent difference in Eif4e or DDX6 (not shown) protein expression.

**Figure 3.**
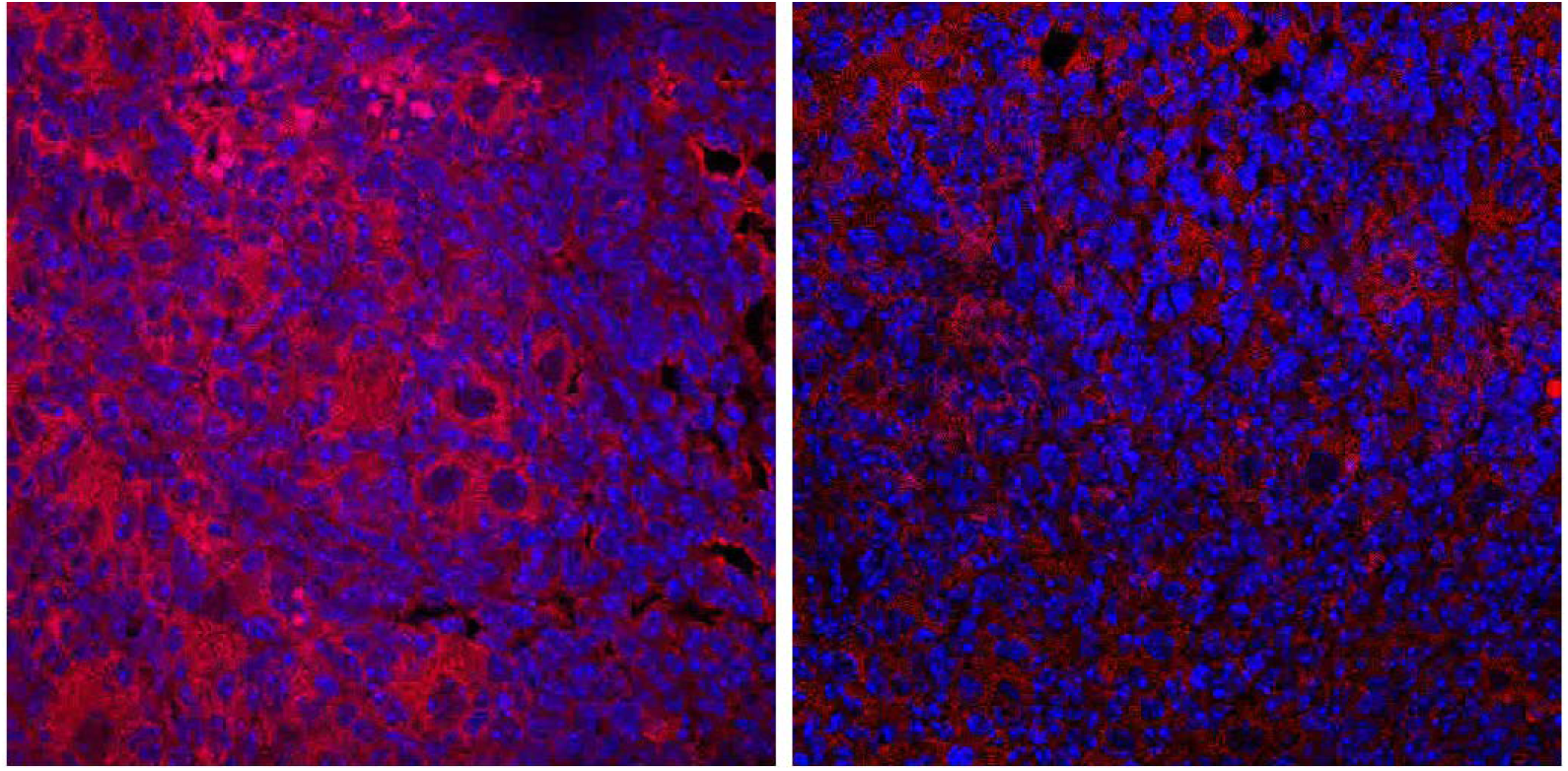
Day 3 Ovaries. Eif4enif1 (red) protein expression in primordial follicles from Day 3 *Eif4enif1^WT/flx^* (left) and *Eif4enif1 ^WT/^*^Δ^ (right) ovaries (60X magnification). Eif4enif1 protein expression is lower day 3 *Eif4enif1 ^WT/^*^Δ^ vs. *Eif4enif1 ^WT/flx^* ovaries.

**Figure 4.**
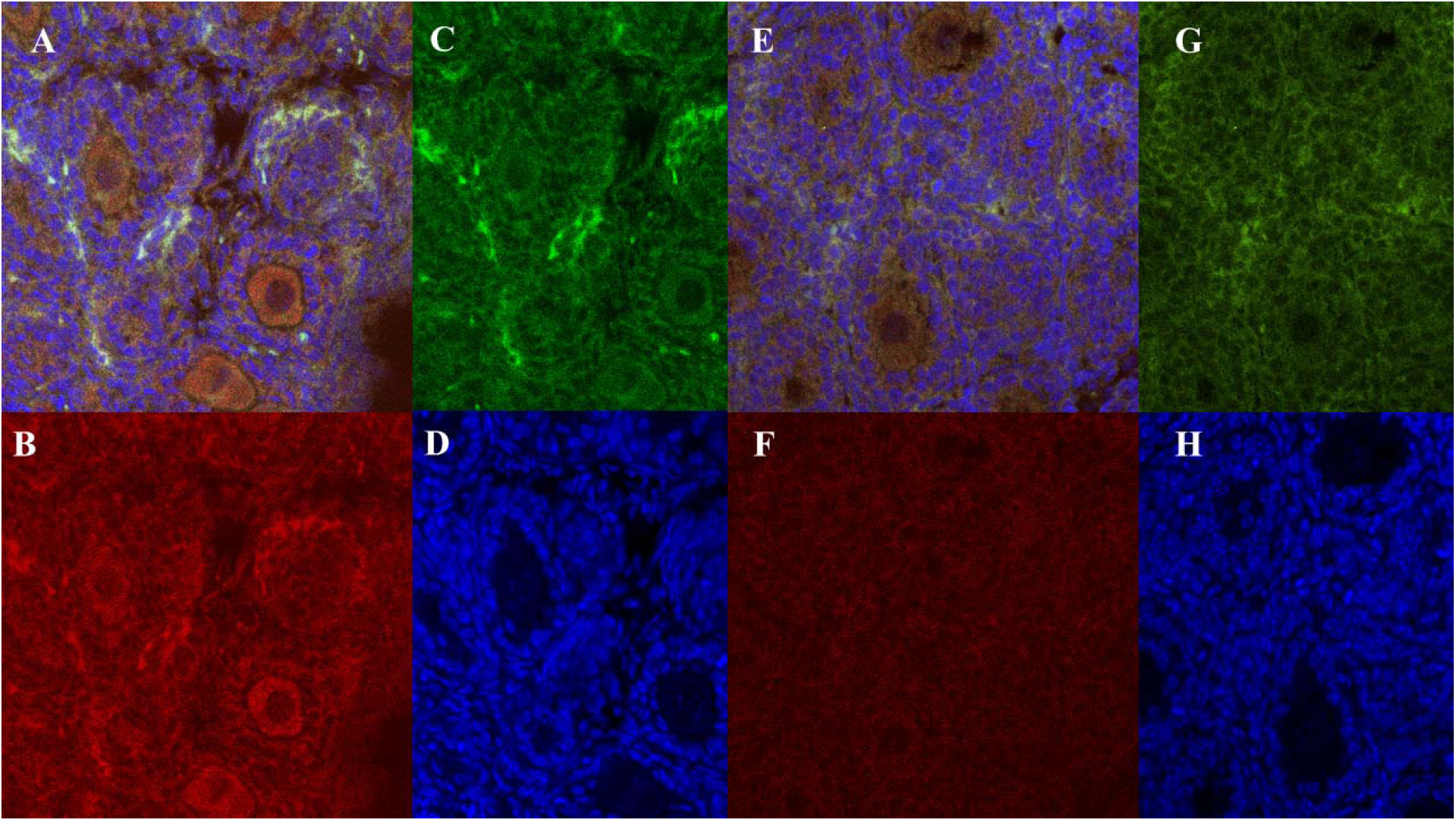
Day 10 Ovaries. A and E) merged, B and F) Eif4enif1 (red), C and G) Eif4e (green) protein expression and D and H) Dapi DNA staining in preantral follicles from day 10 *Eif4enif1^WT/flx^* (A-D) and *Eif4enif1 ^WT/^*^Δ^ (E-H) ovaries (60X magnification). Eif4enif1 protein expression is less intense in *Eif4enif1 ^WT/^*^Δ^ compared to *Eif4enif1 ^WT/flx^* ovaries.

**Figure 5.**
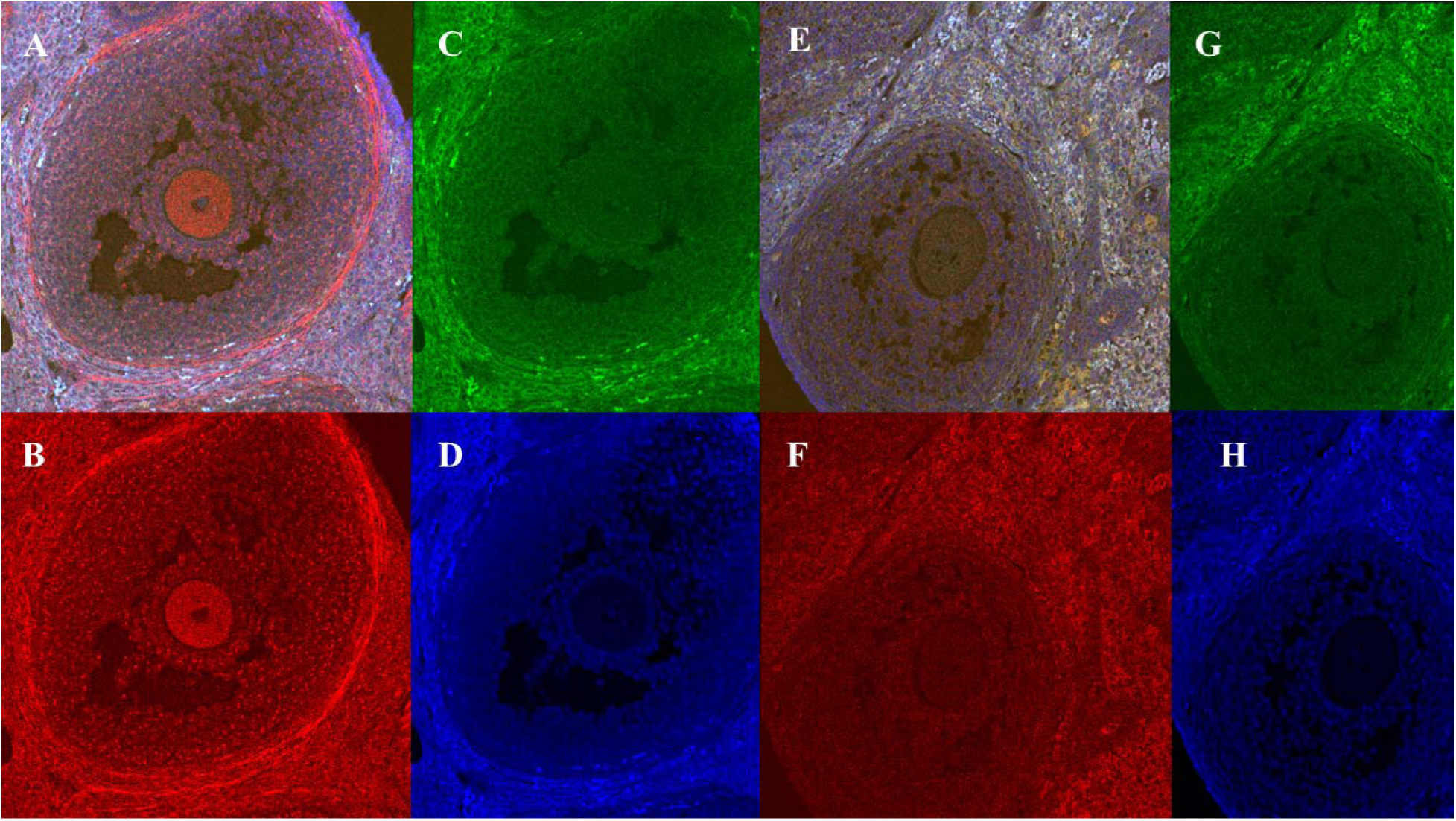
Week 21 Ovary. A and E) Merged, B and F) Eif4enif1 (red), C and G) Eif4e (green) protein expression and D and H) Dapi DNA staining and in an antral follicle from a 21-week old *Eif4enif1^WT/flx^*(A-D) and *Eif4enif1 ^WT/^*^Δ^ (E-H) mouse ovaries (60X magnification).

The relative quantity of both C-terminal and N-terminal Eif4enif1/βactin increased from day 1 to day 10 (p<0.03)(Table 2). The relative quantity of both C-terminal and N-terminal Eif4enif1/βactin was lower in *Eif4enif1 ^WT/^*^Δ^ compared to *Eif4enif1^WT/flx^*(p<0.001). The relative quantity of C-terminal and N-terminal Eif4enif1/βactin was also lower in *Eif4enif1 ^WT/^*^Δ^ compared to *Eif4enif1^WT/flx^* on individual Day 1 and for C-terminal on Day 10 (p<0.05). The relative quantity of *Eif4enif1 ^WT/^*^Δ^ was lower using C-terminal vs. N-terminal antibodies on Day 10 (p<0.05, Table 2).

**Table 2.** Relative protein quantity Eif4enif1/Bactin using N and C terminal antibodies. Protein quantity was lower, overall, in *Eif4enif1 ^WT/^*^Δ^ compared to *Eif4enif1^WT/flx^* (*p*<0.001) for both N terminal and C terminal antibodies (p<0.05).

|  | N Terminal |  | C Terminal |  |  |
| --- | --- | --- | --- | --- | --- |
| Time Point | Het | WT | Het | WT | *p* value<sup>1</sup> |
| Day 1 | 0.022±0.039<sup>a</sup> | 0.55±0.13<sup>b</sup> | 0.016±0.14<sup>c</sup> | 0.31±0.20<sup>d</sup> | 0.001 |
| Day 5 | 0.19±0.16 | 0.73±0.53 | 0.16±0.13 | 0.49±0.33 | 0.01 |
| Day 10 | 0.57±0.36<sup>a,c</sup> | 0.82±0.011<sup>c</sup> | 0.12±0.047<sup>b</sup> | 0.62±0.20<sup>c</sup> | 0.001 |

<sup>1</sup>One-way ANOVA. Different letters indicate differences in Holm-Sidak post hoc testing.

#### IVF

Superovulation experiments demonstrated that there was no difference in oocyte number obtained from *Eif4enif1 ^WT/^*^Δ^ and *Eif4enif1 ^WT/flx^* females (18.7±7.8 vs. 19.2±8.7; p=0.89). There was no difference in proportion of fertilized oocytes that formed blastocysts from *Eif4enif1 ^WT/^*^Δ^ and *Eif4enif1 ^WT/flx^* females fertilized with *Eif4enif1 ^WT/^* ^Δ^ male (0.46±0.22 vs. 0.46±0.27; p=0.96). There was also no difference in the stage at which embryos arrested (p=0.4).

Genotypes followed expected Mendelian ratios overall. However, all GG zygotes arrested at, or before, the 4 to 8 cell stage (Table 3). The genotypes of unfertilized oocytes from *Eif4enif1 ^WT/^*^Δ^ contained equal C and G alleles (19.5±7.8 vs. 14.5±3.5; p=0.5).

**Table 3.** Embryo and blastocyst genotypes from female *Eif4enif1 ^WT/^*^Δ^ *X* male *Eif4enif1 ^WT/^*^Δ^.

| Stage | Genotype | *Eif4enif1*<sup>WT/Δ</sup><br>X *Eif4enif1*<sup>WT/Δ</sup> | Expected | P value |
| --- | --- | --- | --- | --- |
| Blastocyst | CC | 6 | 6.25 | 0.03 |
|  | CG | 19 | 12.5 |  |
|  | GG | 0 | 6.25 |  |
| 4-8 cell | CC | 3 | 4.25 | 0.3 |
|  | CG | 6 | 8.5 |  |
|  | GG | 8 | 4.25 |  |
| 2 cell | CC | 2 | 1.5 | 0.9 |
|  | CG | 2 | 3 |  |
|  | GG | 2 | 1.5 |  |
| Total | CC | 21 | 12 | 0.4 |
|  | CG | 27 | 24 |  |
|  | GG | 10 | 12 |  |

There was no difference in the mean intensity of eIF4ENIF1 immunostaining in the *Eif4enif1 ^WT/^*^Δ^ oocytes compared to *Eif4enif1 ^WT/flx^* (417.4±71.7 vs. 483.2±12.5; p=0.3), with the spindle more intense in the *Eif4enif1 ^WT/flx^* mice (Figure 6A and E). At the 2-cell stage, DDX6 was present in the cytoplasm, similar to Eif4enif1 (Supplementary Figure 5). The mean intensity of immunostaining was punctate at the *Eif4enif1 ^WT/^*^Δ^ fertilized 4-8-cell stage compared to the diffuse staining in 2-cell embryos (Figure 6 B and F), although a punctate pattern began to appear in the *Eif4enif1 ^WT/^*^Δ^ at the 2-cell stage (Figure 6B-D and F-H). The mean intensity of immunostaining decreased from oocyte to blastocyst in both *Eif4enif1 ^WT/^*^Δ^ and *Eif4enif1 ^WT/flx^* (Figure 6A,D and E,H). Of note, one cell remained immunostained at all stages, including in the blastocyst, and this cell was the remnant polar body (Figure 6).

**Figure 6.**
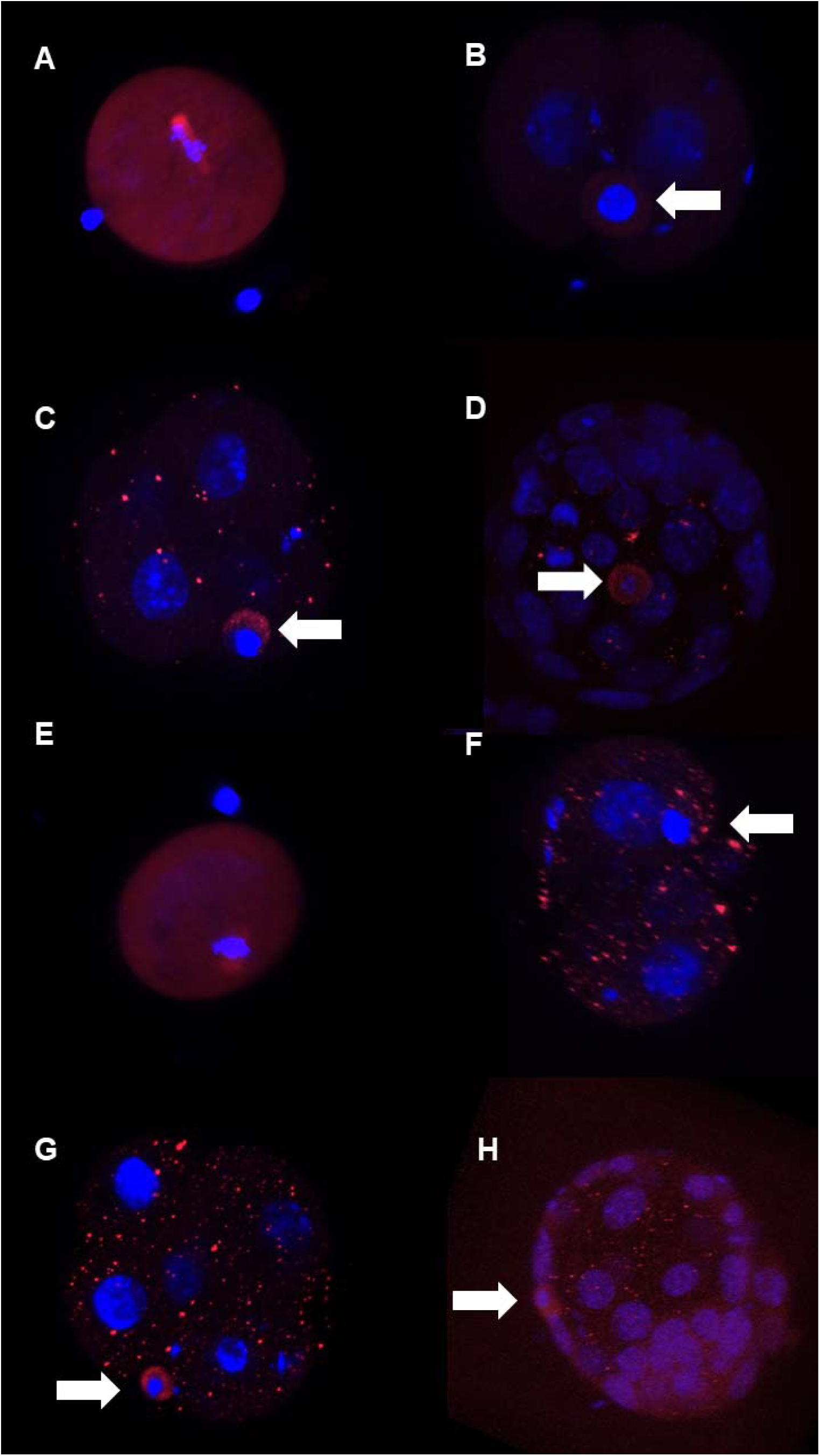
Eif4enif1 protein from meiosis I through blastocyst stage in *Eif4enif1^WT/flx^* (A-D) and *Eif4enif1 ^WT/^*^Δ^ (E-H) mice (red). A and E) Oocytes after meiosis I. Overall Eif4enif1 protein is lower in *Eif4enif1 ^WT/^*^Δ^ (E). In both oocytes, Eif4enif1 is concentrated on the spindle. B and F) Embryos at the 2-cell stage. Note diffuse Eif4enif1 expression in the cytoplasm in the *Eif4enif1^WT/flx^* oocyte (B) but the speckled pattern in the *Eif4enif1 ^WT/^*^Δ^ oocyte (F). The arrow indicates the polar body. C and G) Speckled Eif4enif1 protein expression at the 4-8 cell stage in both genotypes. D and H) Eif4enif1 protein has almost disappeared from the *Eif4enif1^WT/flx^* blastocyst but is still present in a speckled pattern in the *Eif4enif1 ^WT/^*^Δ^ blastocyst. In all images, DNA is stained with Dapi. White arrows indicate the location of the polar body where Eif4enif1 protein is concentrated. All images are shown at 60X magnification.

### Ribo-Seq and Polysome Profiling

When considering translation efficiency for genes with log2≥2 between WT (*Eif4enif1 ^WT/flx^*) and Het (*Eif4enif1 ^WT/^*^Δ^) ribosome protected mRNAs (RPR), the translation efficiency was not different on day 5 (Het vs. WT; 0.82±1.68 vs. 1.05±1.27; p=0.20) or day 22 (0.30±1.29 vs. 1.00±0.80; p=0.47), but was lower on day 10 (0.81±1.45 vs. 3.25±1.95; p<0.001). Using the cutoff of log2≥2, there were more genes that had a negative log2 fold change in RPR for Hets (n=104, 150 and 2 on days 5, 10 and 22, respectively) compared to a positive log2 fold change (n=35, 1 and 1)(Supplementary Tables 1-3), with lower differential reads, overall, on day 22.

The genes with higher ribosome protected mRNA reads in Hets on day 5 included matrix metalloproteinases, cytoskeletal components, aromatase and myeloid and immune cells (Table 4 and Supplementary Table 1). Interestingly, many of these genes had lower reads on day 10 (Table 4 and Supplementary Table 2). On day 22, there were very few ovary specific genes with differential mRNA translation in *Eif4enif1 ^WT/^*^Δ^ mice with the exception of lower *Akr1b7* and *H4c17* (Table 5 and Supplementary Table 3).

**Table 4.**
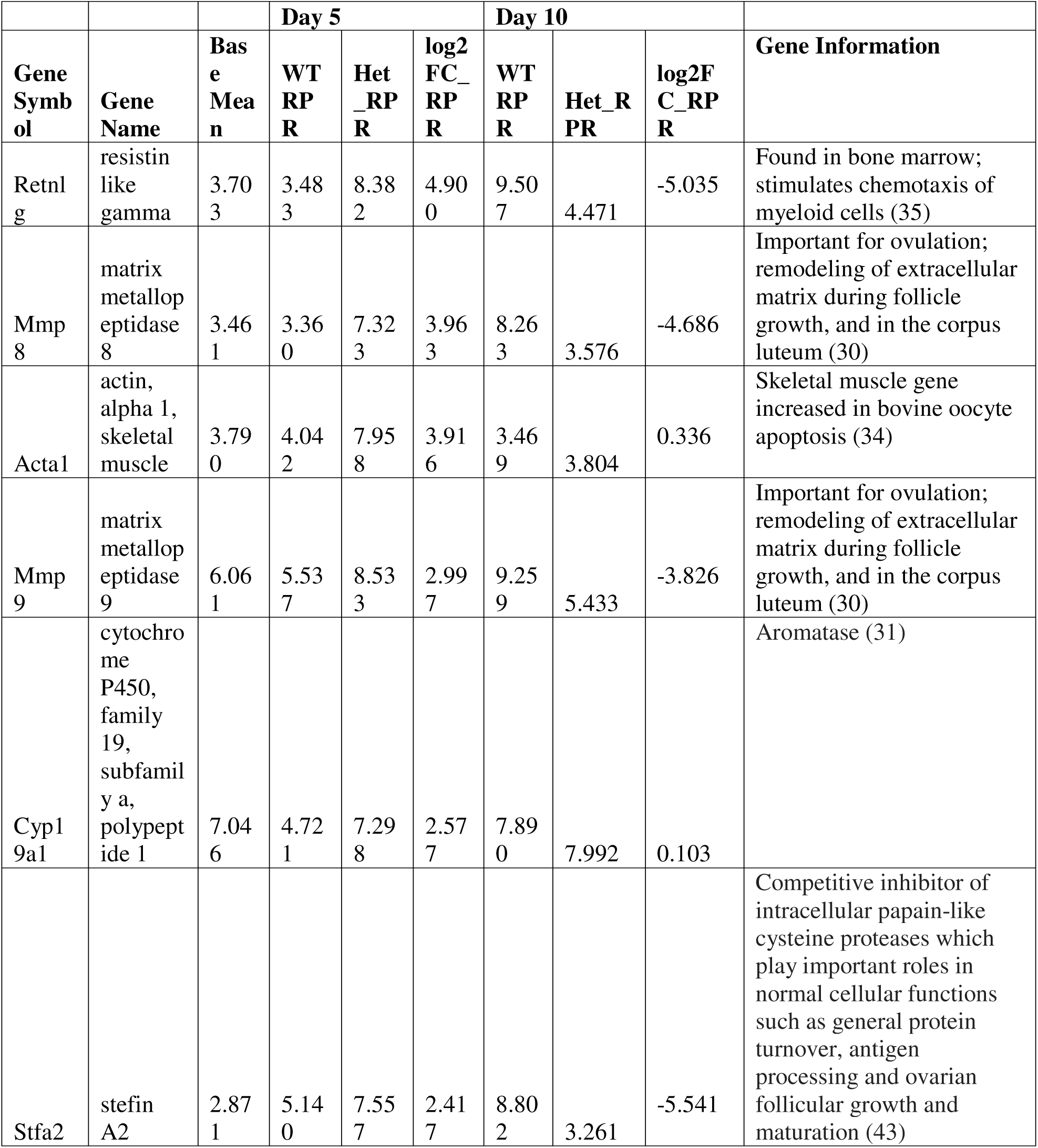
Select genes with a positive or negative log2 fold change (log2FC)≥2 in mRNA translation in Hets compared to WT on days 5 and 10 based on RNASeq of ribosome protected mRNA reads (RPR). Note that many of the genes with mRNA that have increased translation in Hets on day 5 have decreased translation on day 10.

**Table 5.**
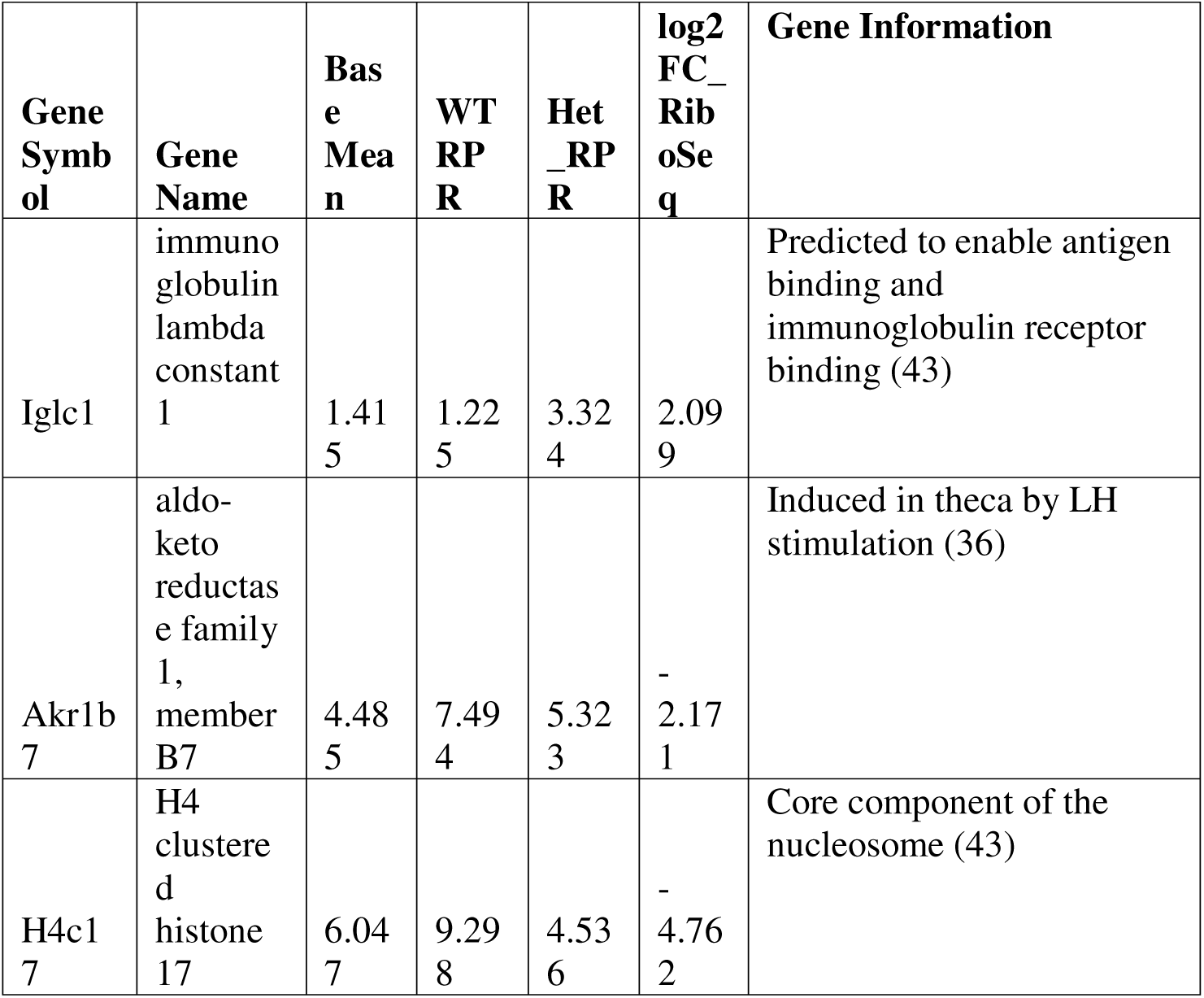
Genes with a positive or negative log2 fold change (log2FC) ≥2 in mRNA translation in Hets compared to WT on day 22 based on RiboSeq of ribosome protected mRNA reads (RPR).

Fast gene set enrichment analysis indicated myogenesis, interferon alpha and gamma response and allograft rejection gene sets were enriched in genes with an increase in translation in Hets on day 5 (log2FC_RPR_5; Supplementary Table 4). The oxidative phosphorylation gene set was enriched for mRNAs with increased translation on day 10. mRNAs with lower reads in Hets on day 10 were mainly in gene sets important for heme metabolism, immune function, Mtorc1 signaling and late estrogen response (log2FC_RPR_10; Supplementary Table 4). The early estrogen response gene set was lower in Hets on day 22 (log2FC_RPR_22; Supplementary Table 4).

Fast gene set enrichment analysis was also performed for oocyte and granulosa cell specific genes from the Ovarian Kaleidoscope to amplify genes important in follicles from the whole ovary samples (19,20). Day 5 was the only day at which there was evidence for an increase in translated mRNA for oocyte specific species in Het *Eif4enif1^WT/^*^Δ^ mice (log2FC_RPR_5; Supplementary Table 5). There were no days in which the translated mRNAs in these genes sets were negatively enriched.

There was no significant difference in *Eif4enif1* total or translated mRNA reads, although the translated mRNA in Hets was approximately 0.87-0.92 of the WT mRNA on days 5-22. The total mRNA reads demonstrated the presence of the c.1283C>G stop-gain mutation in approximately half of the reads. PCR performed on the polysome fraction from day 5 cDNA using primers encompassing *Eif4enif1* exon 9-10 also demonstrated the presence of the stop-gain mutation in actively translated mRNA (Supplementary Figure 6). In the same sample, there was no difference in total polysome mRNA relative units for *Eif4enif1* in *Eif4enif1 ^WT/^*^Δ^ and *Eif4enif1^WT/flx^* (0.002±0.001 vs. 0.0045±0.0007).

### Additional Phenotypes

*Eif4enif1 ^WT/^*^Δ^ had increased skin ulceration, eye opacities, ovarian and tubal cysts and rectal prolapse (Table 6). Reproductive organ abnormalities such as cysts on the ovaries or tubes or blocked tubes appeared to be more common in *Eif4enif1 ^WT/^*^Δ^ mice.

**Table 6.** Additional Phenotypes in in *Eif4enif1 ^WT/^*^Δ^ mice.

| Phenotype | WT (%) | Het (%) | *p* value |
| --- | --- | --- | --- |
| Skin ulceration | 3.1 | 12.4 | <0.001 |
| Eye disease | 0.52 | 8.3 | <0.001 |
| Reproductive Organ | 1.6 | 7.5 | 0.006 |
| Rectal Prolapse | 0 | 3.7 | 0.02 |
| Neurologic Dz | 0 | 2.1 | 0.1 |
| Early Death | 3.1 | 6.6 | 0.4 |

## Discussion

We developed a mouse model with a stop-gain mutation in *Eif4enif1* that recapitulates the *eIF4ENIF1* stop-gain mutation in a family with a dominant form of POI (5). The heterozygous mice have a clear POI phenotype demonstrating decreased follicle number and decreased fertility compared to wild type mice. As in the human family, the mutation can be inherited through females despite decreased fertility. Follicles were lost from the primordial through the preantral stage, starting at day 10.

Homozygous *Eif4enif1* ^Δ*/*Δ^ embryos do not develop beyond the 4 to 8 cell stage, suggesting that Eif4enif1 is critical for embryogenesis. There was evidence for early, increased mRNA translation on day 5, but subsequently decreased translation of the same genes on later days. There was also increased total oocyte and follicle-specific mRNA species with sparing of translated mRNAs in the heterozygotes.

Our data provide the first comprehensive examination of *Eif4enif1* mRNA and protein expression across oocyte and early blastocyst development in mammals. The data demonstrate that *Eif4enif1* mRNA and protein expression begin as early as day 1, appearing in the cytoplasm of primordial and primary follicles. Further, Eif4enif1 increases during the period of rapid oocyte growth from day 1 to day 10 in the cytoplasm of granulosa cells and oocytes. It remains in the cytoplasm of oocytes through meiosis II, expanding previous findings (21). In the fertilized embryo, the protein is diffuse in the cytoplasm at the 2-cell stage, but was more concentrated in the polar body. Eif4enif1 protein disappeared in the transition from the 2-cell embryo to the blastocyst stage, changing from a diffuse to a speckled pattern at the 4-cell stage and then disappearing entirely.

We also demonstrated that Eif4enif1 protein associates with the spindle. Eif4enif1 appeared to bind to the meiotic apparatus in meiosis I oocytes. Of note, a yeast 2 hybrid system previously demonstrated that Eif4enif1 bound to mitotic spindle associated protein p126 (MAP126) or Mastin, which plays a role in spindle formation during mitosis (21). Mutations in the Mastin resulted in male and female infertility related to ovarian hypoplasia (22). The mammalian homolog, astrin, binds to the meiotic spindle after germinal vesicle breakdown, regulating microtubule organization and spindle pole integrity (23). Interestingly, mutations in *Astrin* result in failure of polar body extrusion in mice (23). Taken together, these data suggest a role for Eif4enif1 in meiosis I and II in spindle translation localization or spindle function and formation of the polar body, in addition to a potential role in the growing oocyte.

Eif4enif1 protein is present but decreased in the presence of the *Eif4enif1* stop-gain mutation (*Eif4enif1 ^WT/^*^Δ^) compared to the wild type across oocyte development. The pattern across development is similar to that in the wild type ovaries. Eif4enif1 in the heterozygotes decreases and disappears from the 2-cell embryo to the blastocyst stage, similar to wild type Eif4enif1 protein expression. However, the pattern of Eif4enif1 protein expression became speckled earlier than in the wild type embryos. Of note, there is mRNA in the total and ribosome protected fragments that contain the stop-gain mutation demonstrating that it does not undergo complete nonsense mediated decay. DDX6 binds to the CHD domain, which is not deleted in the stop gain mutation (24). The pattern of higher DDX6 protein expression in the 2-cell *Eif4enif1 ^WT/^*^Δ^ embryos also suggests that a shortened form may remain at this stage of embryo development.

Ovaries with a heterozygous *Eif4enif1* stop-gain mutation (*Eif4enif1 ^WT/^*^Δ^) demonstrate early follicle loss. Oocyte and follicle loss occur across oocyte development from the primordial to preantral stage beginning on day 10. Previous studies used siRNA to block Eif4enif1 during in vitro growth of mouse follicles isolated at day 10-12 (25). siRNA blockade of Eif4enif1 in day 10-12 follicles resulted in failure of nuclear envelope breakdown and failure to resume meiosis. However, some of our *Eif4enif1* stop-gain mutation oocytes were fertilized in the mouse model and the *Eif4enif1* stop gain mutation can be transmitted to progeny in humans through autosomal dominant inheritance (5), suggesting that meiosis can proceed with a reduced amount of Eif4enif1 protein. Therefore, the quantity of Eif4enif1 may be critical for these processes. Taken together with the protein expression patterns, our data suggest that decreased eIF4ENIF1 results in oocyte loss across the period of oocyte growth starting at the primordial stage, with potential effects on meiosis possibly related to decreased full-length protein quantity affecting a subset of oocytes.

We examined apoptosis as a mechanism for follicle loss. There was no evidence of apoptosis on day 5. TUNEL staining, which does not detect oocyte demise, identified granulosa cell apoptosis on days 17 and 22 in *Eif4enif1 ^WT/^*^Δ^ follicles in which the oocyte had pulled away, likely identifying early atretic follicles (26). We also examined oocyte apoptosis through the intrinsic pathway in which a stressor results in release of cytochrome C from the mitochondria, activating Caspase 9 and then caspase 3, which then cleaves poly-ADP ribose polymerase-1 (PARP1)(27). Caspase 3 and PARP1 staining were present in oocytes of the preantral and small antral follicles at week 20, but not in earlier preantral follicles on day 21. The oocyte apoptosis coincides with follicles that were expected to be undergoing resumption of meiosis I (28), which occurs around the time of antrum formation. Therefore, it appears that follicle loss may be related to granulosa cell loss in early stages, whereas oocyte apoptosis may correspond to resumption of meiosis I in some remaining follicles.

Despite follicle and oocyte loss, the number of superovulated oocytes was preserved. The oocyte number was not different between *Eif4enif1^WT/flx^* and *Eif4enif1^WT/^*^Δ^, and the genotype of unfertilized oocytes was not different than the expected Mendelian frequency. These findings suggest movement of follicles from primordial and primary pool maintains antral follicle number at the expense of immature oocytes and follicles. In support, there were very few primary and preantral follicles left in the *Eif4enif1^WT/^*^Δ^ mice at month 10 and these mice had no litters after approximately 6 months as a sign of oocyte depletion.

In addition to follicle loss, our data demonstrate that Eif4enif1 plays an important role in embryogenesis. While the Mendelian expectations were met overall when examining genotypes, there were no homozygous (GG) embryos that progressed beyond the 8 -cell stage after fertilization. The absence of homozygous GG embryos accounts for the lower pup number *Eif4enif1^WT/^*^Δ^. The *Eif4enif1^WT/^*^Δ^ mice appear to lose Eif4enif1 prematurely at the 2-cell stage, which is the last stage containing cytoplasmic Eif4enif1. Maternal mRNAs are eliminated from the fertilized oocyte, with 2300 maternal mRNAs eliminated immediately following fertilization, consistent with a maternal mode of clearance, while almost 500 mRNAs are cleared at the 2-cell stage, consistent with a zygotic mode of regulation (29). Thus, there may be retention or premature loss of maternal mRNAs before the 2-cell stage that results in developmental failure.

Based on the role of *Eif4enif1* in other organisms, we hypothesized that decreased Eif4enif1 resulted in early, increased and uncontrolled mRNA translation, accounting for oocyte and follicle loss (8–11). On day 5, several notable genes affecting follicle growth demonstrated increased translation. These included specific matrix metalloproteinases, which assist tissue breakdown to facilitate follicle growth and ovulation of the oocyte (30). Mmp8 increases to remodel the extracellular matrix during follicle growth and in the corpus luteum (30). Mmp9 expression increases in the later stages of follicle growth, particularly after PMSG stimulation (30). The increased translation of these proteins on day 5 in *Eif4enif1^WT/^*^Δ^ mice suggest that remodeling may occur too early, resulting in abnormal tissue breakdown and dysfunction of normal folliculogenesis.

Cyp19a1 also exhibited increased translation on day 5, indicating potentially increased aromatase at this early stage and the possibility of premature responsiveness to FSH or early development (31). Other genes that were of interest included *Stfa2*, or Stefin A2, which inhibits papain-like cysteine proteases, including cathepsins (32). *Stfa*s are expressed in primordial follicles and inhibit primordial follicle growth by estradiol-mediated inhibition of cathepsins (33), which could be detrimental for normal primordial follicle development. *Acta1* expression is upregulated during apoptosis in bovine oocytes (34), perhaps indicating oocyte distress in the current model. The increase in *Retnlg* might indicate activation and infiltration of inflammatory cells, which would normally be present at the time of ovulation but may also be increased if there is tissue damage from early matrix metalloproteinase action (35).

On days 10 and 22 there were very few genes that demonstrated increased translation. Many of the genes with increased translation on day 5 had decreased translation on day 10, suggesting that the mRNA was depleted early in the *Eif4enif1^WT/^*^Δ^ mice. Day 22 results did not overlap with day 5 or day 10. There was decreased mRNA translation of *Akr1b7,* which is induced in theca by LH stimulation (36) and overall decreased translation of early estrogen response genes. Taken together, there was decreased translation of factors important for steroidogenesis and ovulation on day 22. The difference in the day 22 ovaries likely relates to the mature follicle development and ovulation in mice at their prime stage of fertility.

Whole ovaries were used to examine translated mRNAs and the mRNAs exhibiting increased translation were not oocyte or granulosa cell specific genes, likely related to the relative quantity of tissue types. Therefore, gene set enrichment analysis was performed using the genes specific for oocytes and follicles from the Ovarian Kaleidoscope (20). There was an increase in translated mRNA for oocyte specific species in *Eif4enif1^WT/^*^Δ^ mice. The gene set included *Dmrta1* and *Oosp1* (germ cell differentiation), *Fbxw15* and *Oas1d* (follicle assembly and growth), *Khdc1b* and *Oas1d* (regulation of translation), *Ooep* (regulation of meiotic nuclear division and double strand break repair), *Nlrp4f* and *Ooep* (cystoskeleton and organelle organization), and *Rfpl4* (targets cell cycle proteins for ubiquitin proteasome degradation)(37–42). Based on their function, it is possible that increased translation of these genes disrupted normal oocyte development or that they were alternatively involved in oocyte rescue. Other genes in the set, such as *Prdx1*, are expressed during ovulation and may therefore be prematurely expressed in the *Eif4enif1^WT/^*^Δ^ mice, suggesting mis-timed mRNA translation. Of note, total mRNA for many of the same genes was also increased in the day 5 *Eif4enif1^WT/^*^Δ^ mice. These same genes and others, such as *Obox1* and *Obox2* and *Oosp3* demonstrated increased total mRNA on day 10, without increased translation. These data suggest the possibility that mRNA transcription increased to maintain sufficient translation of mRNA to maintain normal levels of these proteins.

The transgenic mouse model replicates autosomal dominant POI in women with the *eIF4ENIF1* stop gain mutation. The heterozygous stop gain mutation causes follicle loss at the time of rapid oocyte growth after day 5 and during meiosis. Therefore, these studies provide a clear mechanism for dominant inheritance of translation related mutations as a model for POI. Homozygous stop gain mutations are incompatible with normal embryo development. The mechanism for follicle loss appears to be decreased translational repression and mis-timed translation of oocyte developmental proteins. *Eif4enif1* represents one of a growing list of POI genes identified in the pathway of translational control.

## Acknowledgements

We acknowledge Health Science Center Cell Imaging Core at the University of Utah for use of equipment (Nikon), and thank the faculty for their assistance.

