## Supplemental Figures for "Heterozygous *Eif4nif1* Stop Gain Mice Replicate the Primary Ovarian Insufficiency Phenotype in Women"

Abbreviated Title: *Eif4enif1* and POI

Supplementary Figure 1. A) eIF4F Complex and Binding Proteins; eIF4E, the 5’ cap dependent translation initiation factor; eIF4A, the ATP dependent helicase; and PABP, the 3’ poly(a) tail binding protein, all bound to the eIF4G scaffolding protein. eIF4ENIF1 binds to eIF4E at the same site it binds to eIF4G.

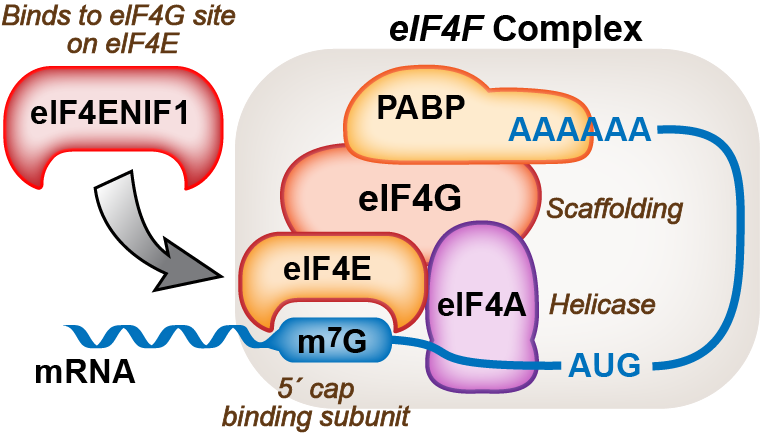

Supplementary Figure 1. B) eIF4E and mRNA Storage or Degradation. eIF4ENIF1 binds to eIF4E at the eIF4G site and to PABP, the poly(a) tail binding protein. The complex then moves to P bodies or germ cell granules where the mRNA is degraded or stored, respectively. Germ cell granules are depicted here.

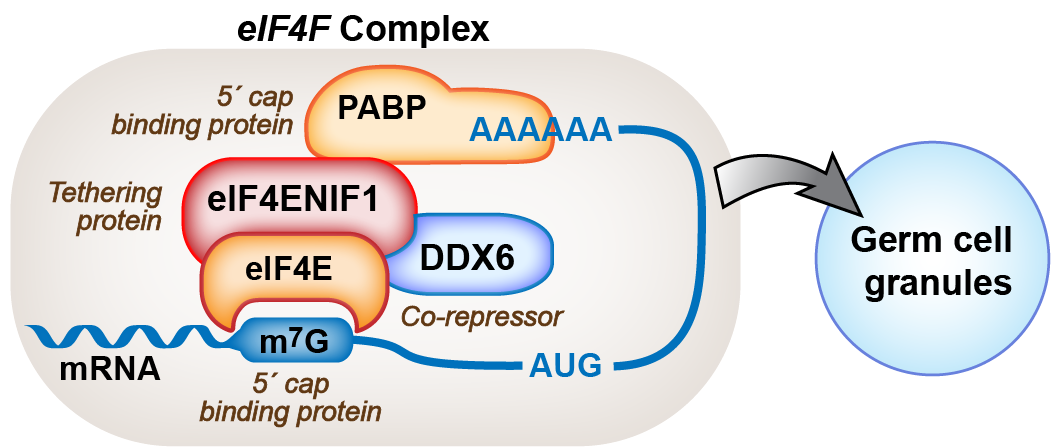

Supplementary Figure 2. Weight in *Eif4enif1^WT/flx^* and *Eif4enif1^WT/Δ^* mice. A) Weight per male mouse in *Eif4enif1^WT/flx^* (black squares) and *Eif4enif1^WT/Δ^* (open circles). B) Weight per female mouse in *Eif4enif1^WT/flx^* (black squares) and *Eif4enif1^WT/Δ^* (open circles). There was no difference in weight in males (p=0.2) or females (p=0.4).
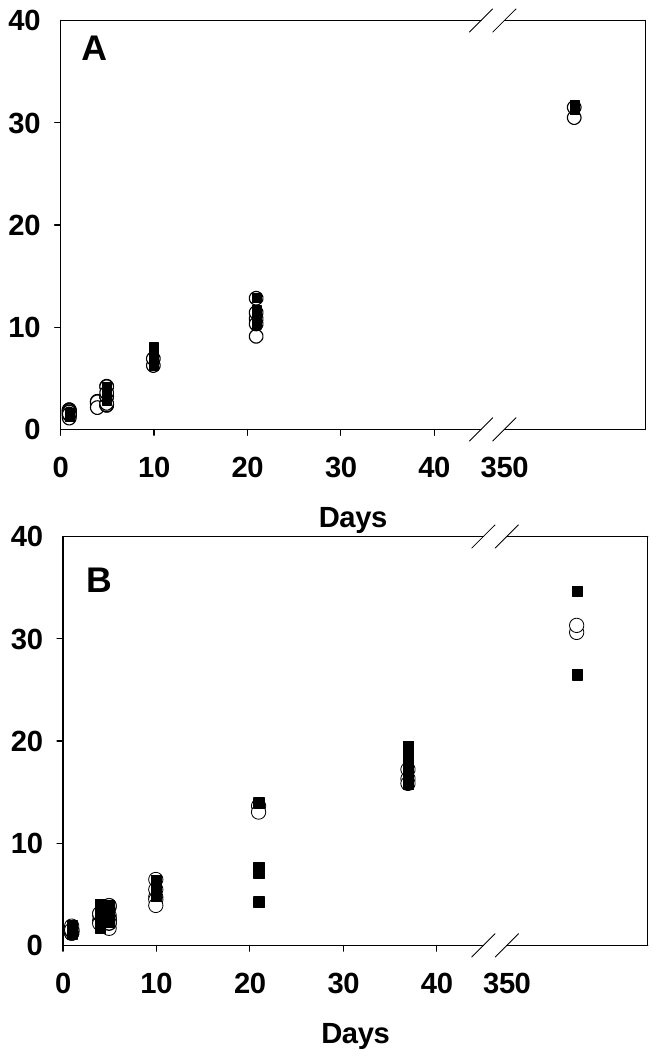

Supplementary Figure 3. Individual testis and ovary volumes. A) Testis volume / Mass in *Eif4enif1^WT/flx^* (black squares) and *Eif4enif1^WT/Δ^* (open circles). B) Ovary volume in *Eif4enif1^WT/flx^* (black squares) and *Eif4enif1^WT/Δ^* (open circles). Testis volume / mass (p=0.4) and ovary volumes (p=0.8) were not different in the two groups.

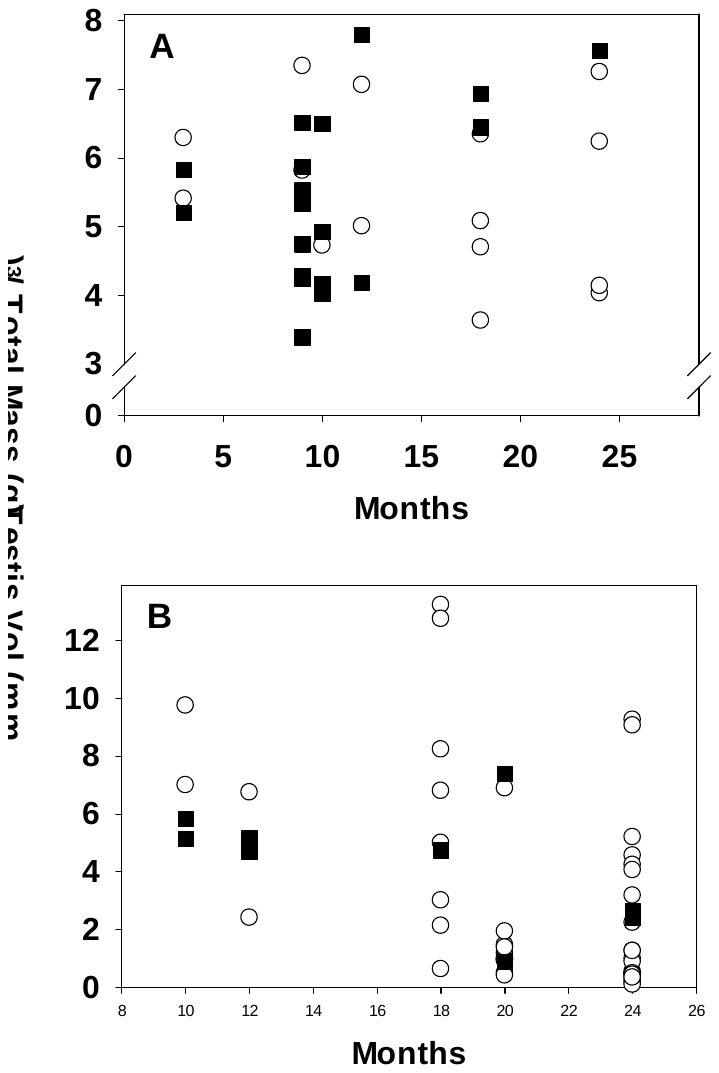

Supplementary Figure 4. Anti-caspase 3 polyclonal antibody detected with DAB staining in an oocyte from a large preantral follicle in a week 22 ovary from an *Eif4enif1^WT^*^/Δ^ mouse. There was no staining demonstrated in *Eif4enif1 ^WT/flx^* mice.

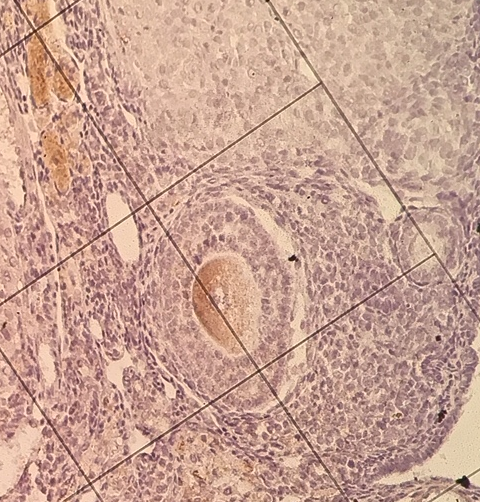

Supplementary Figure 5. Eif4enif1 and DDX6 protein in 2-cell embryos in *Eif4enif1^WT/flx^* (A,C) and *Eif4enif1 ^WT/Δ^* (B,D) mice (red). A and B) Eif4enif1 expression in the cytoplasm in the *Eif4enif1^WT/flx^* (A) oocyte and diffuse/speckled pattern in the *Eif4enif1 ^WT/Δ^* (B). The arrow indicates the polar body. C and D) DDX6 protein concentrated at the dividing cell border in the *Eif4enif1^WT/flx^* blastocyst going from 2- to 4-cell stage, but diffuse in the cytoplasm of the *Eif4enif1 ^WT/Δ^* 2-cell blastocyst. In all images, DNA is stained with Dapi. White arrows indicate the location of the polar body where Eif4enif1 protein is concentrated. All images are shown at 60X magnification.

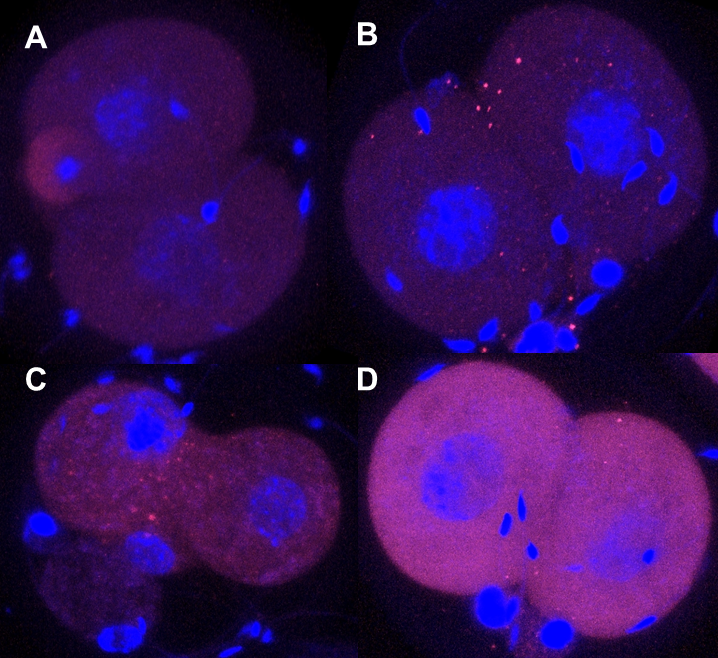

Supplementary Figure 6. Electropherogram and cDNA sequence of mRNA from polysomes of *Eif4enif1^WT/flx^* (top) and *Eif4enif1 ^WT/Δ^* (bottom) mice.

**TTCAGGGG**

**G**

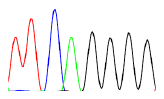

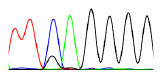
